# QTL mapping of the narrow-branch “Pendula” phenotype in Norway spruce (*Picea abies* L. Karst.)

**DOI:** 10.1101/2022.09.19.508593

**Authors:** Francisco Gil-Muñoz, Carolina Bernhardsson, Sonali Sachin Ranade, Douglas G. Scofield, Pertti O. Pulkkinen, Pär K. Ingvarsson, M. Rosario García-Gil

**Affiliations:** Department of Forest Genetics and Plant Physiology, Umeå Plant Science Centre, Swedish University of Agricultural Science (SLU), 901 83 Umeå, Sweden; Department of Ecology and Genetics, *Evolutionary Biology*, Uppsala University, 752 36 Uppsala, Sweden; Natural Resources Institute Finland (LUKE), 00790 Helsinki, Finland; Department of Plant Biology, Swedish University of Agricultural Sciences (SLU), 756 51 Uppsala, Sweden

## Abstract

Pendula-phenotyped Norway spruce has a potential forestry interest for high density plantations. This phenotype is believed to be caused by a dominant single mutation. Despite the availability of RAPD markers linked to the trait, the nature of the mutation is yet unknown. We performed a Quantitative Trait Loci (QTL) mapping based on two different progenies of F1 crosses between pendula and normal crowned trees using NGS technologies. Approximately 25 % of all gene bearing scaffolds of *Picea abies* genome assembly v1.0 were mapped to 12 linkage groups and a single QTL, positioned near the center of LG VI, was found in both crosses. The closest probe-markers placed on the maps were positioned 0.82 cM and 0.48 cM away from the Pendula marker in two independent pendula-crowned x normal-crowned wildtype crosses, respectively. We have identified genes close to the QTL region with differential mutations on coding regions and discussed their potential role in changing branch architecture.

## Introduction

The molecular control of lateral branching involves phytohormones such as cytokinins, auxin (IAA) and strigolactones (Leyser, 2008). Even though plant architecture-related pathways are fairly well understood in model species (Jiao et al., 2021; Roychoudhry & Kepinski, 2015; Sakai & Haga, 2012; Strohm et al., 2013), the genetic regulation of branching architecture in trees, and especially in conifers, is overall poorly investigated. During the last decade, several studies have been conducted on branching mutants in different angiosperm tree species in order to identify the responsible genes underlying the branching phenotype. The *TAC1* gene is responsible for vertically oriented growth of branches and mutatons in *TAC1* have been shown to result in ‘pillar’ phenotypes in Peach (*Prunus persica*) (Dardick et al., 2013), Plum (*Prunus domestica*) (Hollender, Waite, et al., 2018), *Populus* × *zhaiguanheibaiyang* (Xu et al., 2017) and Black cottonwood (*Populus trichocarpa*) (Fladung, 2021). Similarly, *LAZY1* has been shown to regulate the horizontal orientation of lateral shoots (Xu et al., 2017). Wider branch angles are regulated by *WEEP* gene in peach, causing a more pendulous phenotype (Hollender, Pascal, et al., 2018).

Trees with narrower crowns, either caused by very small or very large branch angles, can potentially be planted closer together and thereby give a possibility to utilize the planting area more efficiently. In Norway spruce (*Picea abies* L Karst.), a branching mutant entitled “Pendula” have been identified and this mutation is characterized by down-oriented lateral branches (Figure 1). Earlier studies have shown that the harvest index of Pendula individuals is higher than wild-type trees; the especially the above-ground one (Pöykkö & Pulkkinen, 1990; Pulkkinen & Poykko, 1990). They also appear to be less sensitive to tree competition than normal-crowned individuals (Gerendiain et al., 2008), which suggests that they can be planted in denser stands. This mutant also appears to segregate in a 1:1 dominant segregation pattern consistent with the control of one or only a few genes (Karki & Tigerstedt, 1985; Lepistö, 1985). However, in order to utilize the Pendula mutant in practical forestry and silviculture, a method for screening the branch phenotype at an early age is necessary. This could be done with a reliable genetic marker. An earlier study aimed at identifying such a marker was conducted by Lehner et al., (1995) using random amplified polymorphic DNA (RAPD) markers to map the Pendula gene using bulked segregant analysis in 43 full-sib progenies from the cross P289 (normal crowned) x E477 (Pendula). One marker, OPH10_720, was found to be linked to the Pendula gene with an estimated recombination frequency of 0.046 (SE = 0.032) (Lehner et al., 1995), but RAPD markers are known to have low reproducibllity, making them unsuitable for large-scale screeings.

**Figure 1.**
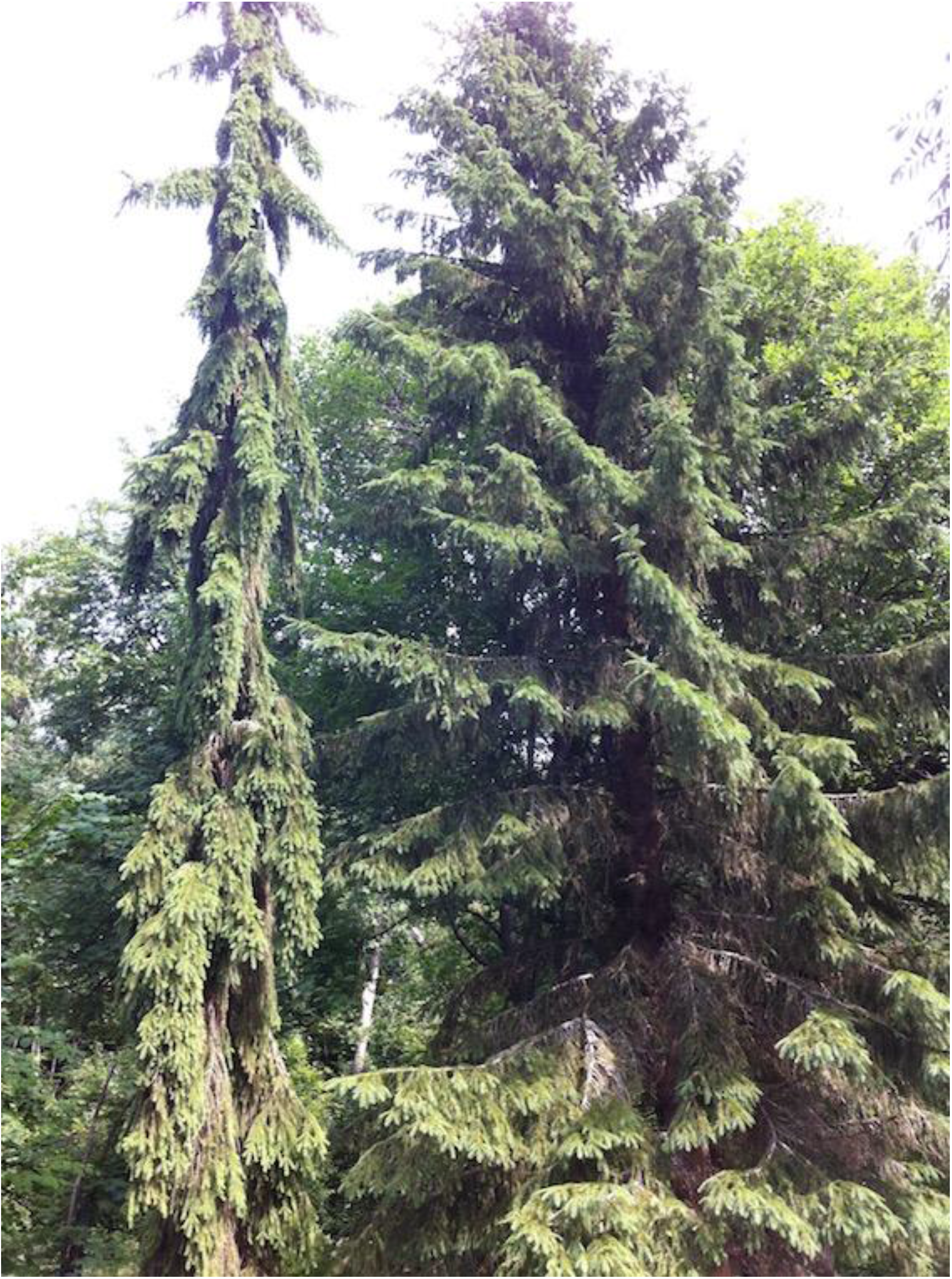
Pendula (left) and regular (right) Norway spruce trees. Photo taken at Arboretum Norr in Umeå, Sweden. Photo taken by Carolina Bernhardsson, summer 2013.

The advent of next-generation sequencing (NGS) technologies and the availability of a draft genome assembly for Norway spruce (Nystedt et al., 2013) have opened up possibilities to explore the genetic architecture underlying the Pendula phenotype by creating dense genetic maps and subsequent Quantitative Trait Loci (QTL) mapping of the phenotype. Here we present four genetic maps, two maternal and two paternal genetic maps, derived from QTL mapping in two independent F1 crosses of a Pendula and normal crowned parent (E477 x K954 and E479 x E2089, respectively).

## Methods & Materials

### DNA extraction and sequence capture

Newly flushed buds were collected from two progeny trials of F_1_ crosses between a Pendula and a normal crowned individual made in the 1980s [E477 (pendula) x K594 and E479 (pendula) x E2089] at a breeding trial run by the Natural Resources Institute Finland (LUKE, formerly METLA) in the spring/summer of 2013. At the same time, branching phenotypes were documented in all offsprings, resulting in 435 and 389 collected and documented progenies for cross E477 x K594 and E479 x E2089, respectievly. The samples were shipped to Umeå University, Sweden, for DNA extraction.

DNA extraction was performed using a Qiagen DNeasy^®^ Plant Mini Kit with approximately 20 ng of freeze-dried tissue as starting material and using the default protocol. Each extracted sample was measured for DNA quality using a Qubit^®^ ds DNA Broad Range (BR) Assay Kit, and all samples had DNA concentrations of ≥ 21.4 ng/μl (mean 66.2 ng/μl) and a total amount of DNA ≥ 2.2μg (mean 7.4 μg). The 828 samples, including samples of the four parents, were sent to RAPiD Genomics^©^ (Gainesville, Florida, USA) in October 2014 for sequence capture using 40,018 capture probes that had been specifically designed to target 26,219 partially validated gene models from the *P. abies* genome assembly (Vidalis et al., 2018). Where possible, probes were designed to flank regions of known contig joins in the v1.0 genome assembly of *P. abies* (Nystedt et al., 2013, for further detail on the probe design, see Vidalis et al., 2018).

The capture data was sequenced by RAPiD Genomics^©^ on an Illumina HiSeq 2000 in either 2×125 or 2×75bp sequencing mode and was delivered in October 2015. Due to the low sequencing depth of the parental samples, these were re-sent for a second round of sequence capture to increase the genotyping call rate of the parents. The raw sequencing reads were mapped against the complete *P. abies* reference genome v.1.0 using BWA-MEM v.0.7.12 (H. Li & Durbin, 2009). The two bam files for each parental sample (caused by the two rounds of sequencing) were merged using Samtools v.1.2 before further processing through the variant calling pipeline. Following read mapping, the BAM files were subsetted to only contain the probe-bearing scaffolds (a total of 24,920 scaffolds) using Samtools v.1.2 (H. Li et al., 2009; H. Li & Durbin, 2009). Duplicates were marked and local realignment around indels was performed using Picard (http://broadinstitute.github.io/picard/) and GATK (https://software.broadinstitute.org/gatk/) (DePristo et al., 2011; McKenna et al., 2010). Genotyping was performed using GATK Haplotypecaller (version 3.4-46), (DePristo et al., 2011; Van der Auwera et al., 2013) with a diploid ploidy setting and gVCF output format. CombineGVCFs was then run on batches of ~200 gVCFs to hierarchically merge them into a single gVCF and a final SNP call was performed using GenotypeGVCFs jointly on the 5 combined gVCF files, using default read mapping filters, a standard minimum confidence threshold for emitting (stand-emit-conf) of 10, and a standard minimum confidence threshold for calling (stand_call_conf) of 20. See Vidalis et al. (2018) for a full description of the pipeline used for calling variants.

### SNP filtering and map creation

The raw VCF-file including the 828 samples was split into each of the crosses separately so that the VCF-file for the E477x K954 cross contained 435 progenies plus parents, and the VCF-file for the E479 x E2089 cross contained 389 progenies plus parents. Samples that showed inconsistency between phenotype labels on the collected tissue bags and in the documentation list (17 and 18 samples to the two crosses, respectively) were removed from the VCF-files before further analysis. The two VCF-files were then filtered so that only bi-allelic SNPs within the extended probe regions (120 ± 100 bp) and without any low-quality tags (QUAL <20) were kept. To increase the chance of capturing the true genotypes, per site sample genotypes were recoded to missing data if they had < 5x coverage or a genotype quality < 10. Principle component analyses (PCAs) were performed on the relatedness estimates from vcftools v. 0.1.12b --relatedness (Danecek et al., 2011), and mislabelled progenies, i.e. progenies not related to both parents, were removed (Supplementary Figure S1). The last pruning step was conducted by removing all SNPs showing >50% missing calls, resulting in a final data set containing 376 samples (including parents) and 333,859 SNPs for the E477 x K954 cross, and 346 samples (including parents) and 317,071 SNPs for the E479 x E2089 cross.

The genotype data were then exported from the VCF-files and all remaining analyses were conducted with R (R core team, 2013). The data sets were thereafter further filtered so that only SNPs where at least one of the parents was heterozygous were kept. Progeny genotype calls were then recoded to missing data if they showed genotypes that were not included in a Punnet square based on parental genotypes, and progenies with > 50% missing calls and SNPs with > 20% missing data were filtered out. A test for segregation distortion was conducted on the remaining SNPs using a chi-square test, and all SNPs with a p-value > 0.005 were kept and considered as informative markers. Each of the informative markers got assigned to the probe region they belonged to and for each probe, only the most informative marker in terms of the least amount of missing data and most balanced segregation pattern was kept for map creation. Finally, the Pendula phenotype was included as a pseudo-genetic marker (Pendula marker) and the data were recoded into BatchMap input format (Schiffthaler et al., 2017). This resulted in 340 F1 progenies (175 pendula and 165 normal crowned) and 9,737 markers for cross E477xK954 and 306 F1 progenies (127 pendula and 179 normal crowned) and 16,687 markers for cross E479xE2089.

Framework genetic maps were then created separately for the two crosses using BatchMap (Schiffthaler et al., 2017), a parallelized version of OneMap (Margarido et al., 2007), using a pseudo test cross strategy. To reduce the number of redundant markers in the map, identical markers (showing no recombination events between them), were grouped into bins and one marker from each bin was used as a bin representative in the map creation. Pairwise estimates of recombination frequencies were calculated between all marker pairs using a LOD score of 14 and a maximum recombination fraction (max.rf) of 0.35. Markers were grouped into linkage groups (LGs) and split into maternal and paternal testcrosses. Each testcross LG was ordered with the “record.parallel()” function using 20 ordering runs over 20 CPU cores. The genetic distance between ordered markers was calculated using the “map.overlapping.batches()” approach with 25 markers overlap between batches, a batch size of ~50 and a ripple window of 11, all parallelized over 2 phase CPU cores and 20 ripple CPU cores. Highly probable miss-ordered markers, i.e. markers showing a recombination fraction distance to the closest neighbor on both sides of > 0.05, were removed and the full testcross LG was recalculated with a new run of “map.overlapping.batches()” and the same settings as previously. Finally, to minimize the effect of genotyping errors on map size, we counted the number of double recombination events in sliding windows of three markers along the testcross LGs and thereafter corrected the genetic distances accordingly.

To anchor the two crosses LGs and give them a common name, the number of shared scaffolds between each LG and a previous haploid consensus map for *P. abies* (Bernhardsson et al., 2019) was used. All LGs were renamed to the haploid consensus maps LG names (Supplementary Figure S2).

### QTL mapping and search for candidate genes

Associations between all markers placed on the genetic maps and the branching phenotype of progenies (pendula or normal crowned) were tested with chi-square tests.

The −log10 (p-value) of the associations were then plotted against the marker position on the testcross to identify the position(s) of any QTLs. To compensate for the lower p-values of double heterozygous markers (segregating in both parents), caused by an extra degree of freedom in the analyses, the p-values for these associations were multiplied by two in the female maps (E477 and E479).

All scaffolds showing marker associations of −log10 (p-value) >40 were considered as candidates for harboring the Pendula locus and all gene models from these scaffolds were extracted and evaluated in Congenie (http://congenie.org) for their gene ontology (GO) and protein family (PFAM) descriptions and compared to the *Arabidopsis thaliana* (*Arabidopsis*) database at http://atgenie.org. Each transcript sequence was also compared to the NCBI blastp database (https://blast.ncbi.nlm.nih.gov) to further analyze the function of the genes (Supplementary File 1).

Candidate genes known to be responsible for branching architecture phenotypes in different angiosperms were positioned on the maps by extracting their corresponding putative conifer sequence ID. This data was obtained either from earlier published articles or, when unknown, by performing the BLAST with known gene model sequences against the Norway spruce draft assembly using blastx at http://congenie.org. The same procedure was performed for genes known to be part of the gravitropism and phototropism pathways in plants (reviewed in Bemer et al., 2017; Hollender, Waite, et al., 2018; Hollender & Dardick, 2015; Jiao et al., 2021; Roychoudhry & Kepinski, 2015; Sakai & Haga, 2012; Strohm et al., 2013) by searching for the genes at atgenie.org and thereafter identifying the corresponding gene models in *P.abies* that belong to the same orthologous gene family and if possible place them on the genetic maps based on scaffold position (Supplementary File 2).

The best match or ortholog for the Norway spruce genes with non-synonymous SNPs was detected in *Arabidopsis* by performing Blastp in PlantGenIE (https://plantgenie.org), TAIR (https://www.arabidopsis.org/index.jsp) and NCBI (https://www.ncbi.nlm.nih.gov/). The domain regions of these genes (Supplementary Figures S3-S8) were confirmed by referring to its best match in *Arabidopsis thaliana* and by performing searches in the Conserved Domain Database (CDD) (Marchler-Bauer et al., 2015), UniProt (Bateman et al., 2021), and referring to the literature (Pang et al., 2014; Wagner et al., 2002).

### Data availability

Raw data is included at https://doi.org/10.5281/zenodo.7093290

## Results & Discussion

Two F1 crosses were used in the QTL mapping of the Pendula phenotype by creating two independent sets of parental genetic linkage maps. These maps were then used to position associations with the qualitative phenotype (pendula or wild-type) using chisquare tests. A total of 19,139 markers from 14,997 gene-bearing scaffolds of the Norway spruce draft assembly v.1.0 (Nystedt et al., 2013) could be placed on the maps. This corresponds to 25.4% of all scaffolds harboring annotated gene models in the v1.0 *P. abies* assembly (14,997 / 58,983 scaffolds), which anchors 9070 high confidence (HC), 6802 medium confidence (MC) and 2042 low confidence (LC) gene models to the genetic maps, corresponding to 26.9% of all partially confirmed gene models in the annotation file (17,914 / 66,632).

The first cross, E477 x K954, contained 340 progenies and 9,714 segregating markers, with 5,525 markers positioned on the female map (E477) and 5,725 markers on the male map (K954). The total size of the parental maps was estimated to 3,585 cM and 3,571 cM, respectively (Table 1). The second cross, E479 x E2089, contained 306 progenies and 16,658 segregating markers, with 9,392 markers positioned on the female map (E479) and 9,786 markers on the male map (E2089). The total size of these parental maps was estimated to 3,393 cM and 3,115 cM, respectively (Table 1). All four parental maps were grouped into 12 LGs each, corresponding to the haploid number of chromosomes in *P. abies* (Sax H & Sax K, 1933).

**Table 1.** Descriptive parameters of the genetic maps.

| LG | E477 (♀ pendula) x K954 (♂ wt) |  |  |  |  | E479 (♀ pendula) x E2089 (♂ wt) |  |  |  |  |
| --- | --- | --- | --- | --- | --- | --- | --- | --- | --- | --- |
|  | E477 markers<br>(bins) | E477<br>size<br>(cM) | K954 markers<br>(bins) | K954<br>size<br>(cM) | Shared<br>markers* | E479 markers<br>(bins) | E479<br>size<br>(cM) | E2089 markers<br>(bins) | E2089<br>size<br>(cM) | Shared<br>markers* |
| **I** | 498 (462) | 364.0 | 575 (522) | 372.8 | 149 | 903 (697) | 309.9 | 900 (710) | 331.2 | 201 |
| **II** | 456 (411) | 282.5 | 500 (448) | 283.6 | 141 | 754 (594) | 252.0 | 829 (642) | 278.7 | 216 |
| **III** | 506 (446) | 288.6 | 417 (374) | 289.9 | 107 | 849 (659) | 298.7 | 818 (650) | 302.4 | 243 |
| **IV** | 462 (403) | 302.2 | 431 (394) | 306.3 | 120 | 750 (577) | 301.8 | 726 (569) | 238.6 | 187 |
| **V** | 466 (416) | 324.1 | 467 (426) | 305.5 | 114 | 785 (600) | 284.1 | 852 (680) | 275.5 | 216 |
| **VI** | 420 (363) | 234.1 | 471 (404) | 243.6 | 108 | 757 (571) | 292.2 | 776 (607) | 211.9 | 209 |
| **VII** | 495 (449) | 341.7 | 513 (451) | 381.7 | 134 | 802 (618) | 295.8 | 930 (700) | 316.7 | 243 |
| **VIII** | 519 (459) | 341.5 | 516 (473) | 330.5 | 145 | 813 (605) | 360.5 | 841 (648) | 246.5 | 184 |
| **IX** | 327 (302) | 254.4 | 440 (397) | 266.2 | 101 | 771 (584) | 285.8 | 816 (604) | 255.0 | 208 |
| **X** | 464 (422) | 275.4 | 459 (417) | 264.4 | 139 | 749 (564) | 243.3 | 748 (569) | 233.2 | 180 |
| **XI** | 440 (368) | 268.5 | 401 (361) | 224.2 | 132 | 596 (440) | 211.6 | 710 (489) | 185.4 | 199 |
| **XII** | 472 (422) | 308.0 | 535 (470) | 302.7 | 146 | 863 (651) | 257.2 | 840 (665) | 240.3 | 234 |
| **Total** | 5,525 (4,923) | 3,585.1 | 5,725 (5,137) | 3,571.5 | 1,536 | 9,392 (7,160) | 3,393.1 | 9,786 (7,533) | 3,115.3 | 2,520 |

The number of unique probe-markers per scaffold that were anchored to any of the two crosses parental maps, ranged between one and six with an average of 1.28 (median 1). 1.4% of all scaffolds (207 out of 14,997) and 5.3% of multi probe-marker scaffolds (207 out of 3,882) have markers that map to different LGs but where the LG grouping is consistent between the two crosses. However, 51 markers (0.27% of all markers and 0.7% of all markers present in both crosses), distributed over 48 different scaffolds, do not show consistent grouping to LGs between the two crosses (supplementary file 3). When comparing the order of shared markers along the LGs between parental maps, the estimated correlations (Kendall’s tau) range between 0.97 and 0.99. Ranges are in line with previously estimated correlations between different maps in *P. abies* (Bernhardsson et al. 2019).

The significance of the associations between marker genotypes and the Pendula phenotype ranged from a −log_10_(p-value) of 0 to 55.1 with a mean of 1.0 and a median of 0.38 for cross E477 x K954. For cross E479 x E2089 the −log_10_(p-value) ranged from 0 to 60.8 with a mean of 1.2 and a median of 0.34. A single large QTL could be detected in each of the crosses, with a peak positioned close to the center of LG VI (Figure 3). The Pendula genetic marker that was added to the linkage maps for reference is positioned in the middle of the QTL in both female maps (black horizontal line for E477 and E479 on LGVI in Figure 3). None of the markers located on any of the other 11 LGs show any evidence of association with the Pendula phenotype. The four parental framework maps show slightly different marker orders (order correlations ranged between 0.97 and 0.99), but the overall genomic patterns are the same (Figure 2). However, this inconsistency in marker order within shorter regions of a genetic map is quite typical and has been seen in several other previously published maps. This is most probably caused by a combination of the heuristic ordering algorithms used for creating dense maps and possible genotyping errors caused by insufficient sequencing depth at some markers (Kelley & Salzberg, 2010; Khan et al., 2012; Salzberg & Yorke, 2005). Since the Pendula phenotype behaves like a qualitative rather than quantitative trait, we chose to included the phenotype as a pseudo-genetic marker in the genetic maps in addition to performing associations against all other markers. The peak of the QTL and position of the Pendula trait ‘marker’ falls at the same location in both maps, which strengthens the robustness of the results. All scaffolds with top associations, −log_10_(p-value) > 40 for markers segregating only in mothers and −log_10_(p-value)*2 >40 for markers segregating in both parents, were investigated for containing candidate genes. In total, 169 probe-markers distributed over 146 scaffolds show high associations. These scaffolds contain 181 annotated gene models of which 131 gene models have orthologous gene family members in *Arabidopsis* (Supplementary File 1).

**Figure 2:**
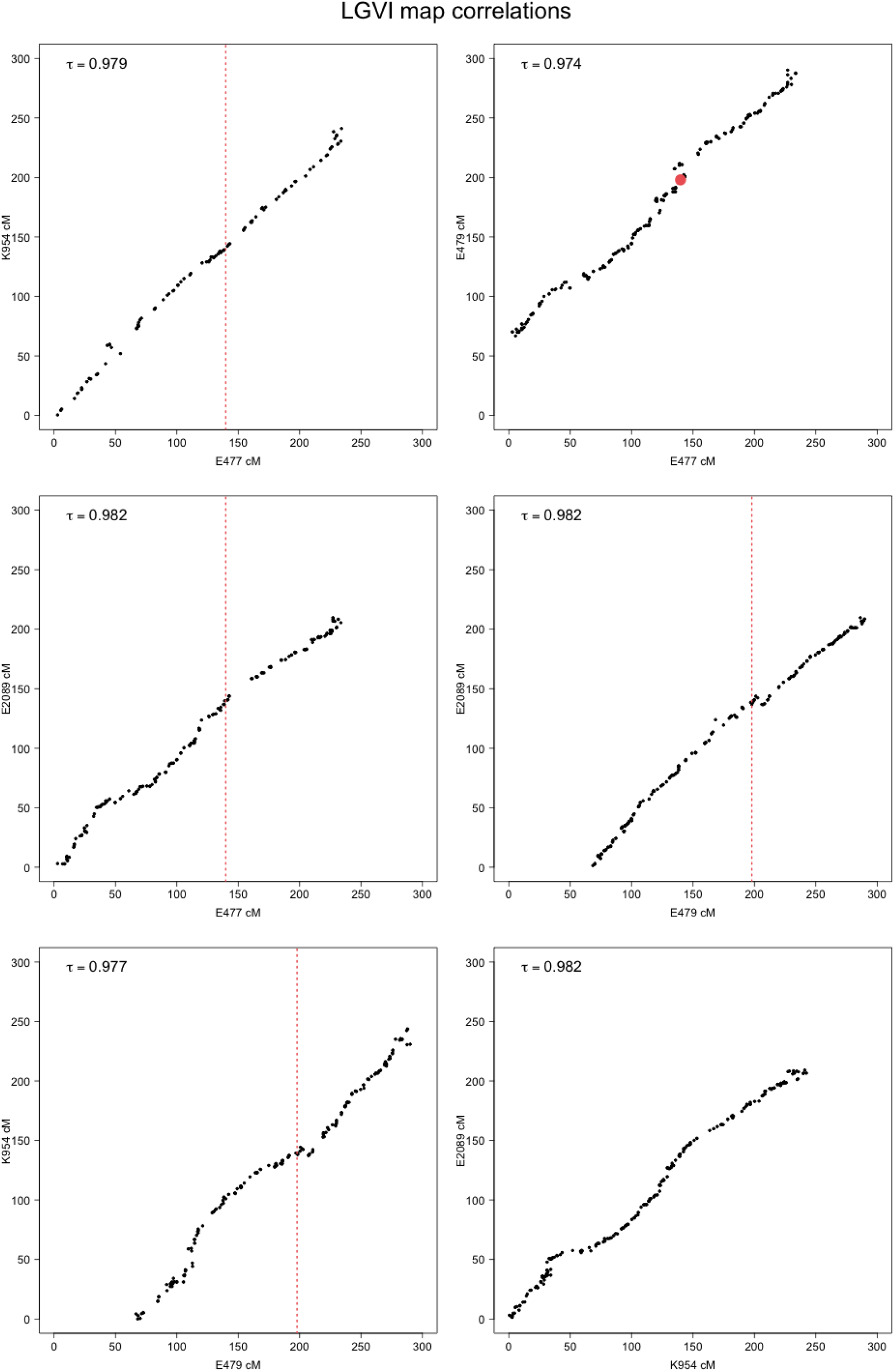
Marker order correlations of LG VI between parental maps. Red dot in top right figure show the position of the pendula marker between E477 and E479. Red dotted lines show the position of the pendula marker when only one of the compared parental maps harbor the marker (E477 or E479). The order correlation, estimated with Kendall’s tau, is shown in the top left corner of each plot.

**Figure 3:**
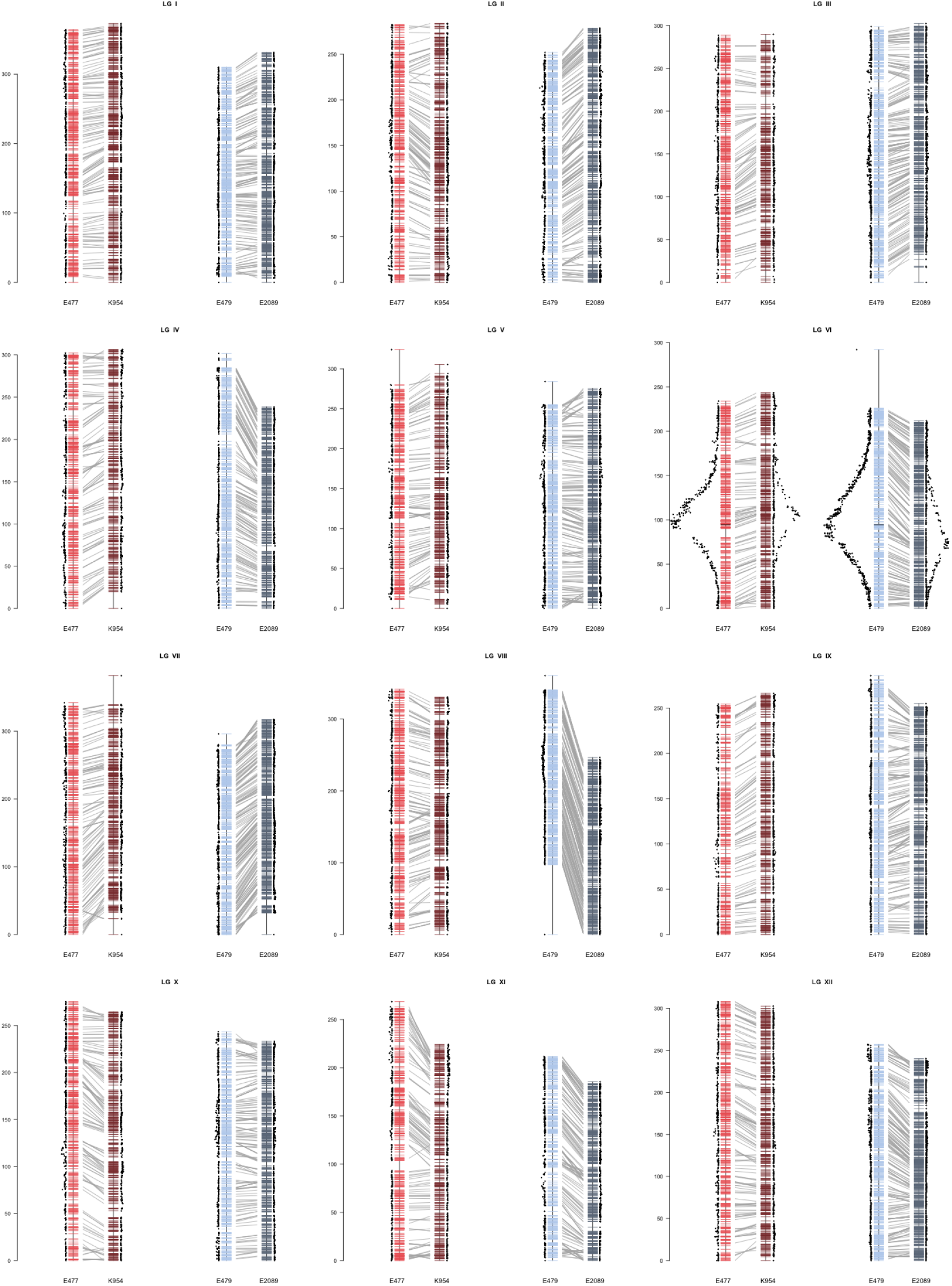
Alignment of the haploid Linkage groups (LG) and significance of the associations between marker genotypes and the Pendula phenotype.

Previously identified genes responsible for branching architecture (Bemer et al., 2017; Hollender, Waite, et al., 2018; Hollender & Dardick, 2015; Jiao et al., 2021; Roychoudhry & Kepinski, 2015; Sakai & Haga, 2012; Strohm et al., 2013) were aligned to the Norway spruce draft assembly and, if found, anchored to the genetic map. A total of 391 gene models were tested, belonging to 28 gene families associated with gravitropism/phototropism (including *WEEP1, WEEP2, LAZY1, TAC1* and *FUL*). Overall, 118 of these gene models (30.2%), from 21 gene families, could be anchored to the genetic maps and are distributed over all 12 LGsand their corresponding putative conifer sequence ID (MA_181281g0010 and MA_10435286g0010 for WEEP, gi|21187158, gi|49450754 for LAZY, gi|69453051 for TAC and no clear sequence for FUL) were identified in the Norway spruce v.1.0 genome assembly. The WEEP genes that have been found to yield a pendulous phenotype in peach (*Prunus persica*) to (Hollender, Waite, et al., 2018) are positioned on LG X (*WEEP1*, MA_181281g0010) and LG V (*WEEP*2, MA_10435286g0010). The IGT gene family, which harbors several genes known to influence plant architecture in angiosperms, including *LAZY*, *TAC* (*Tillar angle control*) and *DRO* (*Deeper rooting*) (Hollender, Waite, et al., 2018; Jiao et al., 2021; Waite & Dardick, 2020), appears to be a single copy gene in *P. abies* as we can only identify a single homolog, MA_39199g0010. This gene is positioned at the distal ends of LG I. 19 of the gene models, from 10 different gene families, are located on LG VI. However, none of the gene models anchored to LG VI are positioned close to the center of the QTL (Supplementary File 2). Even though the list of candidate genes positioned within a 5 cM genetic distance from the Pendula marker is still fairly large (Supplementary Table S1 and S2), we can now rule out most of the earlier known genes involved in tree branching architecture, including the gravitropism and phototropism biosynthesis pathways.

For female E477, 43 probe-markers, from 41 different scaffolds, are positioned within a 5 cM distance from the Pendula marker, with the closest markers occurring at a distance of 0.82 cM (Supplementary Table S1). Seven probe-markers are positioned at this closest distance and one of them, MA_10436629:1, does also show the most significant association for the whole cross (p-value 5.58e-56). This probe-marker is located within the gene model MA_10436629g0010, which transcribes an adenylosuccinate synthase (ADSS) (Supplementary Table S2) and is also present in the cross E479 x E2089, positioned 12.81 cM away from the pendula marker on the E477 map but show a highly significant association (p-value 9.53e-60). This gene class is involved in the *de novo* purine biosynthesis pathway (Stayton et al., 1983), which is a central metabolic function (Smith & Atkins, 2002) and has been described to express at higher levels in the shoot apex vegetative, young leaves and rosette (bar.utoronto.ca/eplant) (Toufighi et al., 2005). To the best of our knowledge, research has not been conducted so far on the effect of alteration in the function of this gene on plant phenotypes, but is known to be the target of a strong herbicide, hydantocydin (Siehl et al., 1996), which again highlights the central role of this enzyme in plant cell metabolism.

For female E479, 31 probe-markers, from 29 different scaffolds, are positioned within a 5 cM distance from the Pendula marker, with two probe-markers positioned at the same position as the Pendula marker (Supplementary Table S2). These two probe-markers are located within the gene models MA_312116g0010, transcribing a lumazine-binding family protein involved in riboflavin synthase/biosynthesis, and MA_10432730g0010, transcribing a P-loop containing nucleoside triphosphate hydrolases superfamily protein. Both of these probe-markers are present as double heterozygotes in the parents (E479 x E2089) but MA_312116:1 shows the most significant p-value of the double heterozygous markers for the cross (p-value 7.17e-30, Table 3). MA_312116:1 is also present as a double heterozygous marker in the E477 x K954 cross, and are there positioned 1.22 cM away from the Pendula marker in the E477 map and shows the strongest association of all double heterozygous markers (p-value 1.14e-31, (Supplementary Table S1). The probe-marker showing the strongest association for cross E479 x E2089 is MA_114136:1 (p-value 1.72e-61) located within the gene model MA_114136g0010, which transcribes a ribosomal protein S26e family protein and is positioned 0.48 cM away from the Pendula marker on the E479 map (Supplementary Table S2).

There are 12 probe-markers significantly associated with the Pendula marker in both crosses (Table 2). Those with the highest p-values are MA_34514 and MA_10429386 located within the gene models MA_34514g0010 and MA_10429386g0010, respectively, which transcribe a membrane trafficking VPS54 family protein. Among these gene models, non-synonymous SNPs were detected in the coding regions of five Norway spruce genes. However, only one of these markers (MA_51707:1) had the same SNP located within the coding region of the gene leading to an aminoacid change in both the crosses. Supplementary Table S3 presents the details of these genes including the alignment information of their corresponding proteins with *Arabidopsis* proteins (Supplementary Figures 3-7) representing the actual alignments performed with MUSCLE (Edgar, 2004). Four genes gave the corresponding BLAST hit in *Arabidopsis* (Supplementary Table S3). The putative *Arabidopsis* orthologues of these genes are involved in various plant processes such as growth, defense and stress response. GLUTATHIONE S-TRANSFERASEs is a huge gene family. One of the members of this family, GLUTATHIONE S-TRANSFERASE U17 (ATGSTU17; AT1G10370) is involved in the modulation of seedling development in *Arabidopsis*. ATGSTU17 participates in the regulation of the architecture of *Arabidopsis* inflorescence by regulating the expression of AtMYB13 gene, which in turn acts at branching points of the inflorescence (Jiang et al., 2010; Kirik et al., 1998). No significant sequence similarity was detected between AT1G10370 and its ortholog in spruce MA_5480022p0010, yet the amino acid Tryptophan (W) where the SNP was located appears to be conserved between both species (Supplementary Figure S7).

**Table 2:** All probe markers positioned within a 5 cM distance from the pendula marker on LGVI for E477 (first) and E479 (second). Marker: name of the probe-marker; Marker position (cM): position on the genetic map; Marker association (segr. Type): chi-square p-value for the association between the probe-marker and the pendula phenotype, segregation type of the probe-marker within brackets; Scaffold: Which scaffold the probe-marker comes from; Scaffold position (bp): Position on the scaffold in bp for which SNP that represents the probe-marker; Gene model: Which gene model the probe-marker belongs to; Arabidopsis ortholog: orthologous gene family in Arabidopsis. Bold names represent markers with the SNP located at the same position in the marker

| Marker | Marker position (cM) | Marker association (segr. type) | Scaffold position (bp) | Gene model and scaffold | Arabidopsis ortholog | Arabidopsis description (synonyms) |
| --- | --- | --- | --- | --- | --- | --- |
| **MA_312116:1** | 197.944<br>138.588 | 7.17e-30 (B3.7)<br>1.14e-31 (B3.7) | **3858**<br>**3858** | MA_312116g0010 | AT2G20690 | lumazine-binding family protein |
| Pendula (E477) | 197.944 | - | - | - | - | - |
| MA_10429386:1 | 198.424<br>137.999 | 7.85e-59 (D1.10)<br>1.77e-50 (D1.10) | 1485<br>1476 | MA_10429386g0010 | AT1G50500 | Membrane trafficking VPS53 family protein (VPS53, HIT1) |
| MA_16252:1 | 198.424<br>137.358 | 1.12e-56 (D1.10)<br>1.66e-50 (D1.10) | 38240<br>38305 | MA_16252g0010 | - | - |
| MA_34514:1 | 198.424<br>137.999 | 1.60e-58 (D1.10)<br>4.23e-54 (D1.10) | 2866<br>2923 | MA_34514g0010 | AT1G50500 | Membrane trafficking VPS53 family protein (VPS53, HIT1) |
| MA_19222:2 | 198.933<br>138.985 | 9.30e-52 (D1.10)<br>1.50e-50 (D1.10) | 1915<br>1856 | MA_19222g0010 | AT5G13650 | Suppressor of variegation 3 (SVR3) |
| MA_45621:1 | 199.211<br>138.985 | 9.73e-26 (B3.7)<br>8.09e-52 (D1.10) | 6130<br>6144 | MA_45621g0010 | AT5G23750 | Remorin family protein |
| Pendula (E479) | 139.811 | - | - | - | - | - |
| **MA_10426685:1** | 200.028<br>141.648 | 3.82e-27 (B3.7)<br>1.53e-53 (D1.10) | **1595**<br>**1595** | MA_10426685g0010 | AT1G74240 | Mitochondrial substrate carrier family protein |
| **MA_51707:1** | 200.618<br>143.118 | 1.72e-57 (D1.10)<br>2.37e-53 (D1.10) | **30163**<br>**30163** | MA_51707g0010 | AT4G04350 | tRNA synthetase class I (I, L, M and V) family protein (EMB2369) |
| **MA_881406:1** | 200.618<br>143.118 | 3.57e-58 (D1.10)<br>1.44e-50 (D1.10) | **7579**<br>**7579** | MA_881406g0010 | AT1G10380 | Putative membrane lipoprotein |
| MA_5480022:1 | 200.618<br>143.118 | 1.57e-57 (D1.10)<br>1.66e-25 (B3.7) | 1587<br>1582 | MA_5480022g0010 | AT2G47730 | Glutathione S-transferase |
| **MA_335371:1** | 200.795<br>143.393 | 1.01e-55 (D1.10)<br>1.79e-53 (D1.10) | **3883**<br>**3883** | MA_335371g0010 | AT1G04960 | Protein of unknown function (DUF1664) |
| MA_873234:1 | 202.325<br>142.561 | 6.64e-56 (D1.10)<br>1.66e-49 (D1.10) | 1588<br>1597 | MA_873234g0010 | AT1G48320 | Thioesterase superfamily protein |

The only marker that presents a common non-synonymous SNP in both crosses is MA_51707.1. Its closest BLAST hit in *Arabidopsis* is AT4G04350 - EMBRYO DEFECTIVE 2369 (EMB2369; tRNA synthetase class I family protein). This gene is highly expressed in young leaves and pedicels (Klepikova atlas). Aminoacyl-transfer RNA synthetases have been identified as key players in translation and they have an early role in protein synthesis (Brandao and Silva-Filho, 2011). It is involved in embryo and plant development (Meinke, 2020). One of the orthologs of this gene was found to be related to the architecture of the embryo and kernel size in maize (X. Li et al., 2022). However, no information regarding phenotype changes in other species is available. We propose that the characterisation of this gene may give insights into tree architecture in spruce.

Since the genetic maps only managed to anchor ~26% of all partially validated gene models in the *P. abies* genome assembly v 1.0 (Nystedt et al. 2013) there is a risk that we missed the causative gene. However, creating a genetic map that anchors all 66,632 validated gene models is not feasible until a less fragmented and more complete genome assembly with additional gene models per scaffold is made available.

## Data availability statement

Raw data is available at https://doi.org/10.5281/zenodo.7093290

## Acknowledgements

This work was supported by grants from the Knut and Alice Wallenberg Foundation and the Swedish Governmental Agency for Innovation Systems (VINNOVA).

## Author contribution

RGG planned the project, POP organized the sample collection, CB extracted DNA, CB and DGS performed alignment and SNP calling, CB and PKI filtered the data and made all maps and QTL analysis, FGM and SSR performed deeper SNP analysis, gene alignments of candidate genes and major modifications to the original draft, CB wrote the first draft, FGM and SRR finalized the draft, all authors commented to the final draft of the manuscript.

## Literature cited

Bateman, A., Martin, M. J., Orchard, S., Magrane, M., Agivetova, R., Ahmad, S., Alpi, E., Bowler-Barnett, E. H., Britto, R., Bursteinas, B., Bye-A-Jee, H., Coetzee, R., Cukura, A., da Silva, A., Denny, P., Dogan, T., Ebenezer, T. G., Fan, J., Castro, L. G.,… Zhang, J. (2021). UniProt: the universal protein knowledgebase in 2021. Nucleic Acids Research, 49(D1), D480–D489. https://doi.org/10.1093/NAR/GKAA1100

Bemer, M., van Mourik, H., Muiño, J. M., Ferrándiz, C., Kaufmann, K., & Angenent, G. C. (2017). FRUITFULL controls SAUR10 expression and regulates Arabidopsis growth and architecture. Journal of Experimental Botany, 68(13), 3391. https://doi.org/10.1093/JXB/ERX184

Bernhardsson, C., Vidalis, A., Wang, X., Scofield, D. G., Schiffthaler, B., Baison, J., Street, N. R., Rosario García-Gil, M., & Ingvarsson, P. K. (2019). An Ultra-Dense Haploid Genetic Map for Evaluating the Highly Fragmented Genome Assembly of Norway Spruce (Picea abies). G3 (Bethesda, Md.), 9(5), 1623–1632. https://doi.org/10.1534/G3.118.200840

Danecek, P., Auton, A., Abecasis, G., Albers, C. A., Banks, E., DePristo, M. A., Handsaker, R. E., Lunter, G., Marth, G. T., Sherry, S. T., McVean, G., & Durbin, R. (2011). The variant call format and VCFtools. Bioinformatics, 27(15). https://doi.org/10.1093/bioinformatics/btr330

Dardick, C., Callahan, A., Horn, R., Ruiz, K. B., Zhebentyayeva, T., Hollender, C., Whitaker, M., Abbott, A., & Scorza, R. (2013). PpeTAC1 promotes the horizontal growth of branches in peach trees and is a member of a functionally conserved gene family found in diverse plants species. Plant Journal, 75(4). https://doi.org/10.1111/tpj.12234

DePristo, M. A., Banks, E., Poplin, R., Garimella, K. V, Maguire, J. R., Hartl, C., Philippakis, A. A., del Angel, G., Rivas, M. A., Hanna, M., McKenna, A., Fennell, T. J., Kernytsky, A. M., Sivachenko, A. Y., Cibulskis, K., Gabriel, S. B., Altshuler, D., & Daly, M. J. (2011). A framework for variation discovery and genotyping using next-generation DNA sequencing data. Nat Genet, 43(5), 491–498. https://doi.org/10.1038/ng.806

Edgar, R. C. (2004). MUSCLE: Multiple sequence alignment with high accuracy and high throughput. Nucleic Acids Research, 32(5). https://doi.org/10.1093/nar/gkh340

Fladung, M. (2021). Targeted crispr/cas9-based knock-out of the rice orthologs tiller angle control 1 (Tac1) in poplar induces erect leaf habit and shoot growth. Forests, 12(12). https://doi.org/10.3390/F12121615

Gerendiain, A. Z., Peltola, H., Pulkkinen, P., Ikonen, V. P., & Jaatinen, R. (2008). Differences in growth and wood properties between narrow and normal crowned types of Norway spruce grown at narrow spacing in Southern Finland. Silva Fennica, 42(3). https://doi.org/10.14214/sf.247

Hollender, C. A., & Dardick, C. (2015). Molecular basis of angiosperm tree architecture. New Phytologist, 206(2). https://doi.org/10.1111/nph.13204

Hollender, C. A., Pascal, T., Tabb, A., Hadiarto, T., Srinivasan, C., Wang, W., Liu, Z., Scorza, R., & Dardick, C. (2018). Loss of a highly conserved sterile alpha motif domain gene (WEEP) results in pendulous branch growth in peach trees. Proceedings of the National Academy of Sciences of the United States of America, 115(20). https://doi.org/10.1073/pnas.1704515115

Hollender, C. A., Waite, J. M., Tabb, A., Raines, D., Chinnithambi, S., & Dardick, C. (2018). Alteration of TAC1 expression in Prunus species leads to pleiotropic shoot phenotypes. Horticulture Research 2018 5:1, 5(1), 1–9. https://doi.org/10.1038/s41438-018-0034-1

Jiang, H. W., Liu, M. J., Chen, I. C., Huang, C. H., Chao, L. Y., & Hsieh, H. L. (2010). A glutathione s-transferase regulated by light and hormones participates in the modulation of arabidopsis seedling development. Plant Physiology, 154(4). https://doi.org/10.1104/pp.110.159152

Jiao, Z., Du, H., Chen, S., Huang, W., & Ge, L. (2021). LAZY Gene Family in Plant Gravitropism. Frontiers in Plant Science, 11, 2096. https://doi.org/10.3389/FPLS.2020.606241/BIBTEX

Karki, L., & Tigerstedt, P. M. A. (1985). Definition and exploitation of forest tree ideotypes in Finland. In Attributes of trees as crop plants / edited by M.G.R. Cannell and J.E. Jackson. [Abbots Ripton, Huntingdon]: Institute of Terrestrial Ecology, 1985.

Kelley, D., & Salzberg, S. (2010). Detection and correction of false segmental duplications caused by genome mis-assembly. Genome Biology, 11.

Khan, M. A., Han, Y., Zhao, Y. F., Troggio, M., & Korban, S. S. (2012). A Multi-Population Consensus Genetic Map Reveals Inconsistent Marker Order among Maps Likely Attributed to Structural Variations in the Apple Genome. PLOS ONE, 7(11), e47864. https://doi.org/10.1371/JOURNAL.PONE.0047864

Kirik, V., Kölle, K., Wohlfarth, T., Miséra, S., & Bäumlein, H. (1998). Ectopic expression of a novel MYB gene modifies the architecture of the Arabidopsis inflorescence. Plant Journal, 13(6). https://doi.org/10.1046/j.1365-313X.1998.00072.x

Lehner, A., Pöykkö, T., Lehner, A., Campbell, M., Wheeler T Pi, N. C., Gliissl, J., Kreike, J., Neale, D. B., & Wheeler, N. C. (1995). Breeding for Crop Trees View project From sustainable design to sustainable implementation-Knowledge value chains for green economy View project Identification of a RAPD marker linked to the pendula gene in Norway spruce (Picea abies (L.) Karst. f. pendula). Theor Appl Genet, 91, 1092–1094. https://doi.org/10.1007/BF00223924

Lepistö, M. (1985). Riippakuusen periytymisestä ja kapealavaisuuden hyväksikäytöstä kuusen jalostukessa. Summary: The inheritance of pendula spruce (Picea abies f. pendula) and utilization of the narrow-crowned type in spruce breeding. Foundation for Forest Tree Breding, Information, 1, 1–6.

Leyser, O. (2008). Strigolactones and Shoot Branching: A New Trick for a Young Dog. In Developmental Cell (Vol. 15, Issue 3). https://doi.org/10.1016/j.devcel.2008.08.008

Li, H., & Durbin, R. (2009). Fast and accurate short read alignment with Burrows-Wheeler transform. Bioinformatics, 25(14), 1754–1760. https://doi.org/10.1093/bioinformatics/btp324

Li, H., Handsaker, B., Wysoker, A., Fennell, T., Ruan, J., Homer, N., Marth, G., Abecasis, G., Durbin, R., & 1000 Genome Project Data Processing Subgroup. (2009). The Sequence Alignment/Map format and SAMtools. Bioinformatics, 25(16), 2078–2079. https://doi.org/10.1093/bioinformatics/btp352

Li, X., Wang, M., Zhang, R., Fang, H., Fu, X., Yang, X., & Li, J. (2022). Genetic architecture of embryo size and related traits in maize. Crop Journal, 10(1). https://doi.org/10.1016/j.cj.2021.03.007

Marchler-Bauer, A., Derbyshire, M. K., Gonzales, N. R., Lu, S., Chitsaz, F., Geer, L. Y., Geer, R. C., He, J., Gwadz, M., Hurwitz, D. I., Lanczycki, C. J., Lu, F., Marchler, G. H., Song, J. S., Thanki, N., Wang, Z., Yamashita, R. A., Zhang, D., Zheng, C., & Bryant, S. H. (2015). CDD: NCBI’s conserved domain database. Nucleic Acids Research, 43(Database issue), D222–D226. https://doi.org/10.1093/NAR/GKU1221

Margarido, G. R. A., Souza, A. P., & Garcia, A. A. F. (2007). OneMap: software for genetic mapping in outcrossing species. Hereditas, 144(3), 78–79. https://doi.org/10.1111/J.2007.0018-0661.02000.X

McKenna, A., Hanna, M., Banks, E., Sivachenko, A., Cibulskis, K., Kernytsky, A., Garimella, K., Altshuler, D., Gabriel, S., Daly, M., & DePristo, M. A. (2010). The Genome Analysis Toolkit: a MapReduce framework for analyzing next-generation DNA sequencing data. Genome Research, 20(9), 1297–1303. https://doi.org/10.1101/gr.107524.110

Nystedt, B., Street, N. R., Wetterbom, A., Zuccolo, A., Lin, Y. C., Scofield, D. G., Vezzi, F., Delhomme, N., Giacomello, S., Alexeyenko, A., Vicedomini, R., Sahlin, K., Sherwood, E., Elfstrand, M., Gramzow, L., Holmberg, K., Hällman, J., Keech, O., Klasson, L.,… Jansson, S. (2013). The Norway spruce genome sequence and conifer genome evolution. Nature 2013 497:7451, 497(7451), 579–584. https://doi.org/10.1038/nature12211

Pang, Y. L. J., Poruri, K., & Martinis, S. A. (2014). tRNA synthetase: TRNA aminoacylation and beyond. Wiley Interdisciplinary Reviews: RNA, 5(4). https://doi.org/10.1002/wrna.1224

Pöykkö, V. T., & Pulkkinen, P. O. (1990). Characteristics of normal-crowned and pendula spruce (Picea abies (L.) Karst.) examined with reference to the definition of a crop tree ideotype. Tree Physiology, 7(1-2-3–4). https://doi.org/10.1093/treephys/7.1-2-3-4.201

Pulkkinen, P., & Poykko, T. (1990). Inherited narrow crown form, harvest index and stem biomass production in Norway spruce, Picea abies. Tree Physiology, 6(4). https://doi.org/10.1093/treephys/6.4.381

R core team. (2013). R Core Team. In R: A Language and Environment for Statistical Computing (Vol. 55).

Roychoudhry, S., & Kepinski, S. (2015). Shoot and root branch growth angle control-the wonderfulness of lateralness. In Current Opinion in Plant Biology (Vol. 23). https://doi.org/10.1016/j.pbi.2014.12.004

Sakai, T., & Haga, K. (2012). Molecular Genetic Analysis of Phototropism in Arabidopsis. Plant and Cell Physiology, 53(9), 1517–1534. https://doi.org/10.1093/PCP/PCS111

Salzberg, S., & Yorke, J. (2005). Beware of mis-assembled genomes. Bioinformatics, 21.

Sax H, & Sax K. (1933). Chromosome number and morphology in the conifers. Journal of the Arnold Arboretum, 14, 356–375.

Schiffthaler, B., Bernhardsson, C., Ingvarsson, P. K., & Street, N. R. (2017). BatchMap: A parallel implementation of the OneMap R package for fast computation of F1 linkage maps in outcrossing species. PLOS ONE, 12(12), e0189256. https://doi.org/10.1371/JOURNAL.PONE.0189256

Siehl, D. L., Subramanian, M. v, Walters, E. W., Lee, S.-F., Anderson, R. J., & Toschi, A. G. (1996). Adenylosuccinate Synthetase: Site of Action of Hydantocidin, a Microbial Phytotoxin. Plant Physiol, 110, 753–758.

Smith, P. M. C., & Atkins, C. A. (2002). Purine biosynthesis. Big in cell division, even bigger in nitrogen assimilation. Plant Physiology, 128(3), 793–802. https://doi.org/10.1104/PP.010912

Stayton, M. M., Rudolph, F. B., & Fromm, H. J. (1983). Regulation, Genetics, and Properties of Adenylosuccinate Synthetase: A Review. Current Topics in Cellular Regulation, 22(C), 103–141. https://doi.org/10.1016/B978-0-12-152822-5.50008-7

Strohm, A., Baldwin, K., & Masson, P. H. (2013). Gravitropism in Arabidopsis thaliana. Brenner’s Encyclopedia of Genetics: Second Edition, 358–361. https://doi.org/10.1016/B978-0-12-374984-0.00662-8

Toufighi, K., Brady, S. M., Austin, R., Ly, E., & Provart, N. J. (2005). The Botany Array Resource: e-Northerns, Expression Angling, and promoter analyses. The Plant Journal, 43(1), 153–163. https://doi.org/10.1111/J.1365-313X.2005.02437.X

Van der Auwera, G. A., Carneiro, M. O., Hartl, C., Poplin, R., del Angel, G., Levy-Moonshine, A., Jordan, T., Shakir, K., Roazen, D., Thibault, J., Banks, E., Garimella, K. V., Altshuler, D., Gabriel, S., DePristo, M. A., Auwera, G. A., Carneiro, M. O., Hartl, C., Poplin, R.,… DePristo, M. A. (2013). From FastQ Data to High-Confidence Variant Calls: The Genome Analysis Toolkit Best Practices Pipeline. In Current Protocols in Bioinformatics (pp. 11.10.1–11.10.33). John Wiley & Sons, Inc. https://doi.org/10.1002/0471250953.bi1110s43

Vidalis, A., Scofield, D. G., Neves, L. G., Bernhardsson, C., García-Gil, M. R., & Ingvarsson, P. K. (2018). Design and evaluation of a large sequence-capture probe set and associated SNPs for diploid and haploid samples of Norway spruce (Picea abies). BioRxiv, 291716. https://doi.org/10.1101/291716

Wagner, U., Edwards, R., Dixon, D. P., & Mauch, F. (2002). Probing the diversity of the Arabidopsis glutathione S-transferase gene family. Plant Molecular Biology, 49(5).https://doi.org/10.1023/A:1015557300450

Waite, J. M., & Dardick, C. (2020). IGT/LAZY family genes are differentially influenced by light signals and collectively required for light-induced changes to branch angle. BioRxiv, 2020.07.15.205625. https://doi.org/10.1101/2020.07.15.205625

Xu, D., Qi, X., Li, J., Han, X., Wang, J., Jiang, Y., Tian, Y., & Wang, Y. (2017). PzTAC and PzLAZY from a narrow-crown poplar contribute to regulation of branch angles. Plant Physiology and Biochemistry: PPB, 118, 571. https://doi.org/10.1016/J.PLAPHY.2017.07.011

